# Pathogenic potential prediction of *Vibrio parahaemolyticus* by using pangenome data with high performance machine learning algorithms

**DOI:** 10.1101/2025.04.08.647818

**Authors:** Zhuosheng Liu, Zhuoheng Li, Jiawei Zhang, C Titus Brown, Luxin Wang

## Abstract

The presence of *Vibrio parahaemolyticus* (*Vp*) at various stages of seafood production has adversely affected public health and threatened the sustainability of the industry. To address the critical public health threats posed by this prevalent seafood-borne pathogen, this research applied advanced machine learning (ML) and deep learning (DL) algorithms to predict the pathogenic potential of *Vp* using pangenome data. Utilizing comprehensive pangenomic assemblies and sophisticated ML/DL models, this study achieved robust and precise pathogenic potential prediction of *Vp* based on source attribution, which provides a novel reliable diagnostic tool facilitating conventional serotyping and virulence gene combination approaches. Based on results, non-core regions in *Vp* pangenome exhibited useful signals ML models can utilize in pathogenic potential determination process. Tree-based ensemble learning methods (Random Forest and Gradient Boosting Trees) have shown the distinguished performance with AUC score 0.97 based on selected pangenome matrix. Furthermore, Convolutional Neural Network successfully predicted the pathogenic potential of isolates with slightly better performance with AUC score 0.98 based on full pangenome. Critical biological insights revealing critical pathogenic potential-associated genes were retrieved from established ML/DL models: the gene feature weight analysis from Random Forest revealed the importance of accessory genes during *Vp* evolution (similarly highlighted by Gram-cam analysis of Convolutional Neural Network), which provided potential guidance for future research direction.

## Introduction

*Vibrio parahaemolyticus* (*Vp*) is a Gram-negative, rod-shaped and halophilic pathogenic bacterium. The wide presence of *Vp* in different production stages of seafood has generated negative impacts on both public health and the seafood industry worldwide, which results in 80,000 illnesses annually in the United States (1). The Economic Research Service of the United States Department of Agriculture (USDA ERS) estimated that the total cost of illnesses caused by *Vp* was $40 million in 2013 (1, 2). These evidences underscore the imperative needs to control *Vp* at different food production stages.

The pathogenic potential of *Vp* is multifactorial, encompassing various virulence factors such as the thermostable direct hemolysin (TDH), TDH-related hemolysin (TRH), and two type III secretion systems (T3SS1 and T3SS2) (3, 4). While these well-known pathogenic markers are being used as screening tools for identifying pathogenic *Vp*, intensive literature has indicated the high genetic diversity of *Vp* isolates, even for the same serotypes (5–7). The availability of next generation sequencing (NGS) revolutionize the diagnostics and characterization of infectious diseases and pathogens; it overcomes limitations associated with the conventional culture-based methods as well as molecular techniques such as the polymerase chain reaction (8–10). Whole genome sequencing can also reveal emerging virulence factors and elucidate strain-level virulence difference (11). Currently, there are 2069 completed *Vp* whole genome sequencing data available in NCBI public databases. However, there are several challenges associated with the utilization and interpretation of these data. First of all, given the volume, complexity and clinical uncertainty of data generated in whole-genome sequencing, expertise is needed for the interpretation of identified genetic variation. Secondly, since *Vp* is not a model microorganism, a large number of genes in *Vp* pangenome still lack accurate and informative annotations based on Kyoto Encyclopedia of Genes and Genomes and Gene Ontology databases. The correlation between these unidentified or putative functional genes and actual pathogenic virulence has not been investigated either (12). How to better utilize the WGS data for better and more accurate detection, differentiation, and characterization of pathogenic *Vp* warrant prompt research efforts.

In recent years, the utilization of machine learning (ML) and deep learning (DL) methods that leverage whole genome sequencing (WGS) data has garnered considerable attention and interest (13). ML, characterized by their ability to learn useful patterns and make predictions from data, have emerged as powerful tools to analyze and interpret WGS data (14). Learning algorithms can identify complex relationships and patterns within the genome, enabling the discovery of genetic variations, functional elements, and associations with virulence serotypes (15). As a special family of ML algorithms, deep learning (DL) can automatically learn useful features from input data without human involvement, which has been proven to have great performance in various tasks such as image recognition and natural language processing (16). It is promising to bring these powerful DL tools onto the field of microbial food safety. The integration of ML and DL methods with WGS data has the potential to enhance our understanding of the pathogenicity basis of bacterial pathogens, facilitate later stage decision making process, and contribute to advancements in final pathogen control.

Therefore, the goal of this study is to address the urgent need for effective control measures against *Vp* by fast and accurate fundamental pathogenic potential prediction using ML and DL algorithms. This study aims to leverage ML/DL methods and WGS data to advance the characterization *Vp* pathogenicity, thus benefiting the risk management of Vp in food production chain.

## Materials and Methods

### Learning model input dataset preparation

Complete genome assemblies of *Vp* were downloaded from NCBI genebank (total accessed number: 2069). For the consistency of genome annotation, the downloaded complete genome assemblies were re-annotated using Prokka (17). Genome labeling was web-scripted based on genome assembly metadata. Isolation source, host information, and sample description information were used to determine the classification of clinical and non-clinical isolates in the genome file. A total of 1981 isolates were used in ML tasks, of which 872 were clinical isolates and 1109 were non-clinical isolates. Pangenome was later constructed based on reannotated genome assemblies uisng Roary (18). The pangenome constructed by Roary has 101422 unique genes based on 1980 *Vp* strains. It is hypothesized that potential virulence-associated biomarkers distribute more in the whole pangenome than the core genome.

### Unsupervised clustering algorithm implementation

Unsupervised clustering was first applied to check if pangenome exhibits distinct grouping patterns that can steadily recognized by learning algorithms. Four distinct types were considered: core genome, soft-core genome, core genome (core with soft-score) plus shell genome, and full pan-genome. Initially, Principal Component Analysis (PCA) was employed to visualize the distribution of data points within these matrices. This step aided in understanding the inherent structure and variance present in the datasets. For clustering, we implemented both Gaussian Mixture Models (GMM) (19, 20). The GMM approach was selected for its ability to model data as a mixture of several Gaussian distributions, allowing for the accommodation of clusters of different shapes and sizes. In parallel, the K-MODES algorithm was utilized, because k-modes algorithm was specifically selected due to its suitability for clustering boolean matrices (binary data, represented as 1 or 0). These two methods provided a view of the clustering patterns within the pan-genome data relying on gene presence/absence signals without supervised guidance.

### Machine learning model establishment

To achieve optimal binary classification results based on features from different layers in pangenome, this study experimented different machine learning algorithms. As a single-parametric classification method, KNN operates on a simple yet effective principle: the classification of any given data point is predominantly influenced by the majority class among its ‘k’ closest neighbors in the dataset (21). The hyperparameter k was set to 5 after testing various values of k, ensuring a balanced blend of precision and generalization. The Support Vector Machine (SVM) was first selected not solely for its versatility but also for its proven track record in adeptly handling diverse datasets (22). To further enhance its capabilities, it was fine-tuned with the Radial Basis Function (RBF) kernel—a choice inspired by the kernel’s prowess in capturing intricate non-linear relationships. Two pivotal parameters were adjusted to optimize the SVM’s performance: the regularization parameter, C, was calibrated to 0.88, striking a delicate balance between achieving accurate classification of training examples and maximizing the decision function’s margin; and the gamma parameter, a hallmark of the RBF kernel, was set at a value of 0.005, thereby determining the extent to which a single training example could influence the classification. Ensemble tree-based models including Random forest and Gradient Boosting Trees were fitted into pangenome features (23, 24). These two tree-based ensemble learning methods were chosen for its inherent robustness and ability to navigate the complexities of multifaceted datasets. The Random Forest was configured to harness the collective power of 100 decision trees. Each of these trees was granted the liberty to grow to a staggering depth of 1000, ensuring a thorough and exhaustive exploration of all conceivable data structures. Naive Bayes, a probabilistic classification technique grounded in Bayes’ theorem, was also included in this study. This algorithm is particularly renowned for its simplicity and efficiency in handling high-dimensional data, especially under the assumption of feature independence (25). To make all model performance comparable using the same data split and ensure reproducibility in the results (10 cross-fold validation), a steadfast random state of 42 was instated.

### Deep learning model establishment

For the DL models of this study, two distinct architectures were employed to predict the pathogenicity of *Vp*. The Multi-Layer Perceptron (MLP) model comprises four fully connected layers. The first layer takes an input of a specified size and outputs to 1400 nodes. Subsequent layers reduce the dimensionality step-wise, with the second layer maps 1400 nodes to 512, the third maps from 512 to 128, and the final layer maps 128 nodes to the output size. Between each fully connected layer, dropout regularization was applied to prevent overfitting. The dropout probability is set at 0.6. The model uses the ReLU activation function for intermediate layers, promoting non-linearity in the learning process.

The Convolutional Neural Network begins with a standard convolutional layer followed by batch normalization and ReLU activation, and a max-pooling layer. The convolutional layers in the model utilize 3×3 filters. It then sequentially connects two residual blocks, increasing the depth and complexity of the model. Residual Block consists of two convolutional layers each followed by batch normalization (26). Post-convolutional layers, the model includes a series of fully connected layers with dropout for regularization. The network uses ReLU activation functions for these layers. The final fully connected layer maps to the number of classes, and a softmax function is applied for multi-class classification. Both models were trained for 100 epochs with a batch size of 256. These configurations aimed to strike a balance between model complexity and computational efficiency while capturing intricate genomic patterns for accurate pathogenicity prediction. DL models were trained on a single NVIDIA A100 GPU provided by the Davis Data Lab at the University of California, Davis.

### Interpretations of random forest and convolutional neural network

To identify important gene features contributing to the pathogenicity prediction of *Vp*, we used two complementary approaches: feature importance analysis with Random Forest and gradient weighted class activation mapping (Grad-CAM) from a Convolutional Neural Network (CNN). The RF model assigned weights to individual genes, ranking them based on their contribution to the model’s predictions and highlighting key features critical for distinguishing pathogenic and non-pathogenic samples. Simultaneously, Grad-CAM was employed to visualize CNN predictions by generating heatmaps that pinpoint regions of the input matrix most influential in the model’s decision-making (27). Together, these methods provided robust evidence for the roles of specific genes in pathogenicity, combining statistical rigor with model interpretability to support the identification of key gene targets for further investigation.

### Model evaluation metrics

To evaluate the classification performance of ML and DL prediction models built in this study, the following metrics were selected: accuracy was calculated to measure the overall correctness of the model, which was calculated as the ratio of correct predictions to the total number of samples. Precision measured the model’s ability from the aspect of prediction, calculated as the ratio of true positive predictions to the total positive predictions made. Recall measured the model’s ability from the aspect of ground truth positives, which was calculated as the ratio of true positives to the sum of true positives and false negatives. The F1-Score, the harmonic mean of precision and recall, was used as a balanced metric considering both the precision and the recall of the classifier. Additionally, to provide a comprehensive view on model performance on slightly unbalanced input data, the Area Under the Curve (AUC) of the Receiver Operating Characteristic curve was applied as a performance metric for classification models. The ROC curve plotted the true positive rate against the false positive rate at various threshold settings. A steeper ROC curve led to a larger AUC value, indicating a better model performance. AUC values closer to 1.0 suggested excellent model discrimination.

## Results

### Overview of *Vp* Pangenome

Pangenome was constructed based on the complete assemblies of 1981 *Vp* strains and was classified into four pangenome categories, core, softcore, shell and cloud, consistent with findings reported by (28). The distribution and quantity of each categorical pangenome calculated by Roary were captured in Fig 1a and Table 1. The pangenome construction analysis of *Vp* revealed 1,161 core genes present in all strains, serving as the foundation of the genetic identity of *Vp*. Additionally, 1,003 soft core genes were identified in 95% to 99% of strains. The analysis also uncovered 3,229 shell genes present in 15% to 95% of strains, likely associated with environmental adaptations and niche colonization. Remarkably, the vast genetic diversity of *Vp* is highlighted by 96,029 cloud genes found in fewer than 15% of strains.

**Figure 1:**
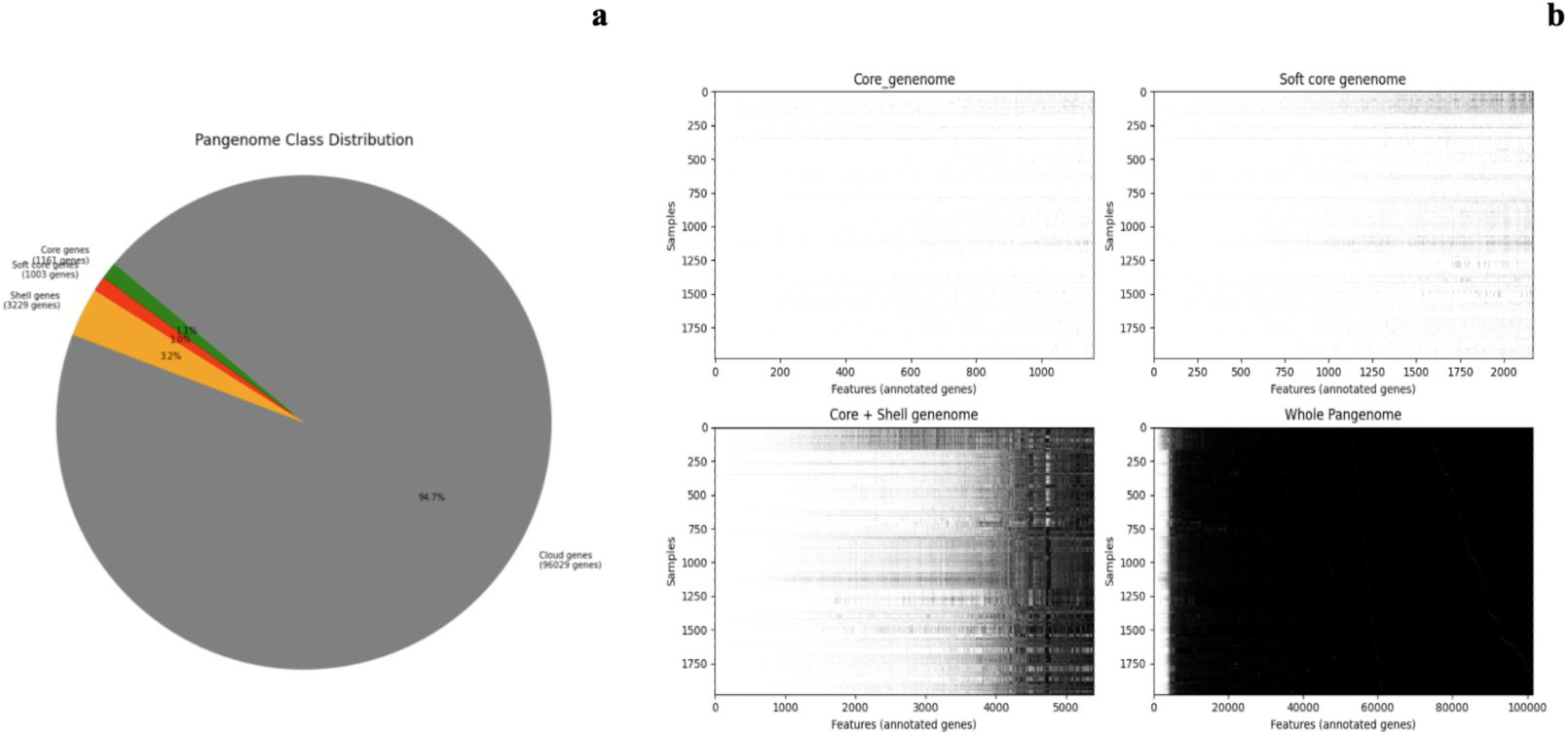
Overview of constructed pangenome of *Vibrio parahaemolyticus* based on complete assemblies of 1980 strains. (a) The pangenome class distribution shows the proportion of core, soft core, shell, and cloud genes. Core genes (1,161 genes) are present in all strains, indicating their essential role in *Vp* biology. Soft core genes (1,003 genes) are found in 95-99% of strains, suggesting strain-specific adaptations. Shell genes (3,229 genes) occur in 15-95% of strains, likely linked to environmental adaptation and niche colonization. Cloud genes (96,029 genes) appear in fewer than 15% of strains, potentially contributing to unique traits, pathogenicity, or resistance mechanisms. (b) Heatmap representations of gene presence-absence across strains in various genome components: Core genome, Soft core genome, Core + Shell genome, and Whole Pangenome. Each row represents a strain, and each column represents a feature (annotated gene), with darker shading indicating gene presence. The Whole Pangenome reveals the high genetic diversity in *Vp*.

**Table 1:** Distribution of genes across strains by pangenome classes.

| **Gene Category** | **Strains (%)** | **Number of Genes** | **Sparsity (%)** |
| --- | --- | --- | --- |
| Core Genes | $99 \leq \text{strains} \leq 100$ | 1,161 | 0.31% |
| Soft Core Genes | $95 \leq \text{strains} < 99$ | 1,003 | 2.80% |
| Shell Genes | $15 \leq \text{strains} < 95$ | 3,229 | 39.76% |
| Cloud Genes | $0 \leq \text{strains} < 15$ | 96,029 | 99.43% |
| **Total Genes** | $0 \leq \text{strains} \leq 100$ | 101,422 | 95.44% |

The heatmaps in Fig 1b demonstrated the gradual increase of sparsity from core genome to the core with shell genome in the gene feature matrix. A high sparsity of matrix was observed in whole pangenome based gene feature matrix (Table 1). The high sparsity on pangenome matrix was caused by the presence of large portion of cloud genes, which were defined as the genes from very few genomes (29).

### Unsupervised clustering is unable to distinguish between clinical and non-clinical *Vp* isolates

The left column of the Fig 2 represents the actual source distribution of *Vp* strains, which was visualized using PCA and categorized as clinical (orange dots) and non-clinical (blue dots), across three different pangenome matrices: core genome (top row), soft core genome (middle row), and core and shell genome (bottom row). These plots serve as the baseline to compare the clustering results from unsupervised clustering algorithms. Notably, as the number of gene features increased from the core genome through the soft core genome and finally to the core and shell genome, the separation between clinical and non-clinical strains became more distinctive based on PCA visualization (Fig2). In the core genome, the two classes showed considerable overlap, which indicated the limited separability offered by 1161 core pangenome genes. This was due to the fact that genes in the core genome were shared among 99% of *Vp*, and therefore *Vp* strains with core pangenome lacked the variability needed for effectively distinguishing different clusters. With the inclusion of additional gene features less commonly shared among strains to the core genome, the two classes (clinical vs non-clinical) began to exhibit better separation, though some overlap persisted. In the core and shell genome, the separation was even more distinct, suggesting that the incorporation of accessory genes improved the differentiation between the two classes. This trend further underscored the importance of a comprehensive pangenome representation in capturing the genomic variation associated with clinical and non-clinical strains.

**Figure 2:**
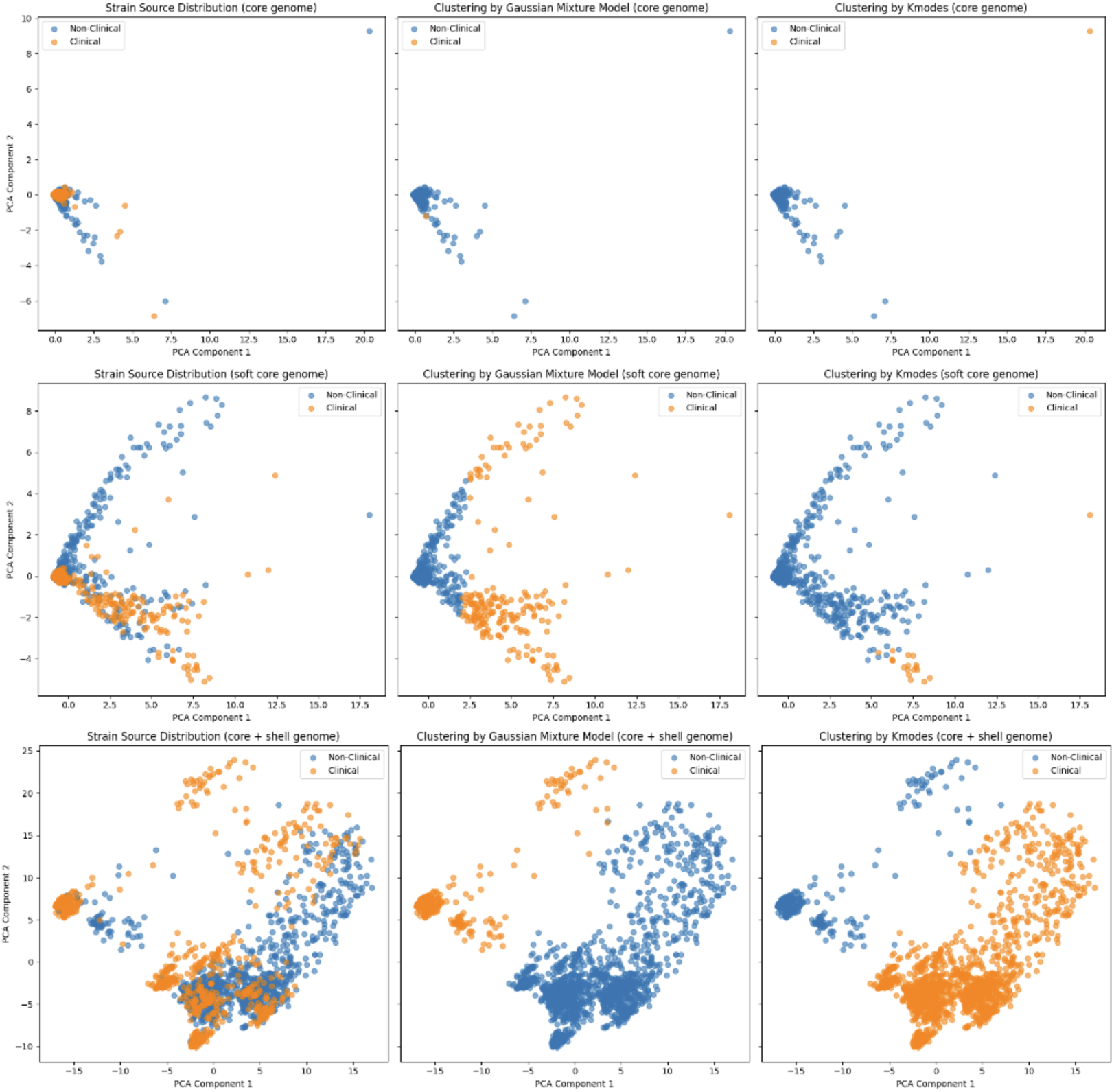
The PCA plots show the clustering of strains based on their source (clinical vs. non-clinical) using PCA components across three genomic categories: core genome (top row), soft core genome (middle row), and combined core + shell genome (bottom row). The left column represents the actual source distribution of strains, while the middle and right columns show clustering results using Gaussian Mixture Models and K-modes clustering, respectively. Orange dots represent clinical strains, and blue dots represent non-clinical strains. The differences in clustering patterns reveal insights into how genomic features influence the differentiation of clinical and non-clinical strains.

Unsupervised clustering algorithms (GMM and K-modes) on three different pangenome matrices (core, soft core, core and shell) were then examined. Table 2 highlights the evaluation of unsupervised clustering algorithms (GMM and K-modes) across different genome types and results demonstrated the unsuitability of unsupervised clustering for classifying clinical versus non-clinical *Vp* strains. For the core genome, both models failed to cluster pathogenic potential of strains based on core-gene presence signals, achieving zero across all metrics. Performance for the soft core genome was similarly poor, with GMM showing minimal effectiveness (F1-Score = 0.21) and K-modes exhibiting high precision (0.95) but virtually no recall (0.02), resulting in an F1-Score of 0.04. On the core and shell genome, both models achieved slightly better results; GMM had moderate precision (0.72) but low recall (0.26), while K-modes delivered higher recall (0.74) but low precision (0.39), leading to an F1-Score of 0.51, the highest among all tested configurations. The whole pangenome matrix was excluded from the unsupervised clustering because the algorithms failed to converge due to its high dimensionality and sparsity, which posed significant challenges to the optimization processes of unsupervised models. Similar poor performance of unsupervised clustering based on PCA-processed Shiga-toxin producing *Escherichia.coli* was reported before (15).

**Table 2:** Unsurpervised clustering of clinical and nonclinical strains based on different genome classes.

| **Genome Type** | **Model** | **Accuracy** | **Precision** | **Recall** | **F1-Score** |
| --- | --- | --- | --- | --- | --- |
| Pangenome | GMM | N/A<sup>a</sup> | N/A<sup>a</sup> | N/A<sup>a</sup> | N/A<sup>a</sup> |
|  | Kmodes | N/A<sup>a</sup> | N/A<sup>a</sup> | N/A<sup>a</sup> | N/A<sup>a</sup> |
| Core + Shell Genome | GMM | 0.63 | 0.72 | 0.26 | 0.38 |
|  | Kmodes | 0.37 | 0.39 | 0.74 | 0.51 |
| Soft Core Genome | GMM | 0.57 | 0.54 | 0.13 | 0.21 |
|  | Kmodes | 0.57 | 0.95 | 0.02 | 0.04 |
| Core Genome | GMM | 0.57 | 0.0 | 0.0 | 0.0 |
|  | Kmodes | 0.57 | 0.0 | 0.0 | 0.0 |

<sup>a</sup> N/A indicates that the model failed to converge during training.

### Performance of ML models initially limited to core- and soft- pangenome was improved on soft and shell and whole-pangenome

Given the failure of unsupervised clustering methods to classify clinical versus non-clinical strains using pangenome signals, supervised classification approaches were subsequently investigated. Five different classic machine learning models (SVM, random forest, K-nearest neighbors, Gradient boosting trees, Naive Bayes) were first fit into four different pangenome matrices (core, soft core, core and shell, pangenome), of which performance evaluation was reported in Fig 3 and Table 3.

**Figure 3:**
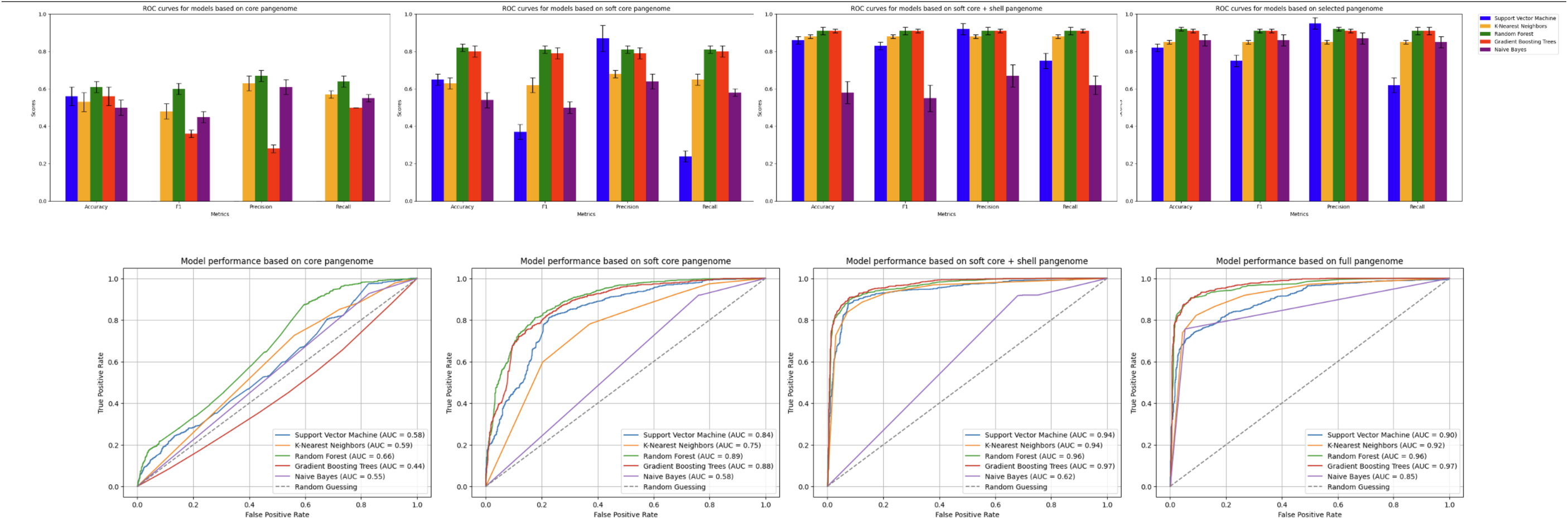
Performance metrics (accuracy, F1-score, precision, and recall; top) and ROC curves (bottom) for ML models (Random Forest, SVM, KNN, XGBoost, Naïve Bayes) trained on core genome, soft core genome, core + shell genome, and whole pangenome datasets. Bar plots show mean scores with error bars representing standard deviation. ROC curves include AUC values for each model, demonstrating superior performance of Gradient Boosting Trees and Random Forest, particularly on larger genomic datasets. Naïve Bayes consistently underperforms across all datasets.

**Table 3:**
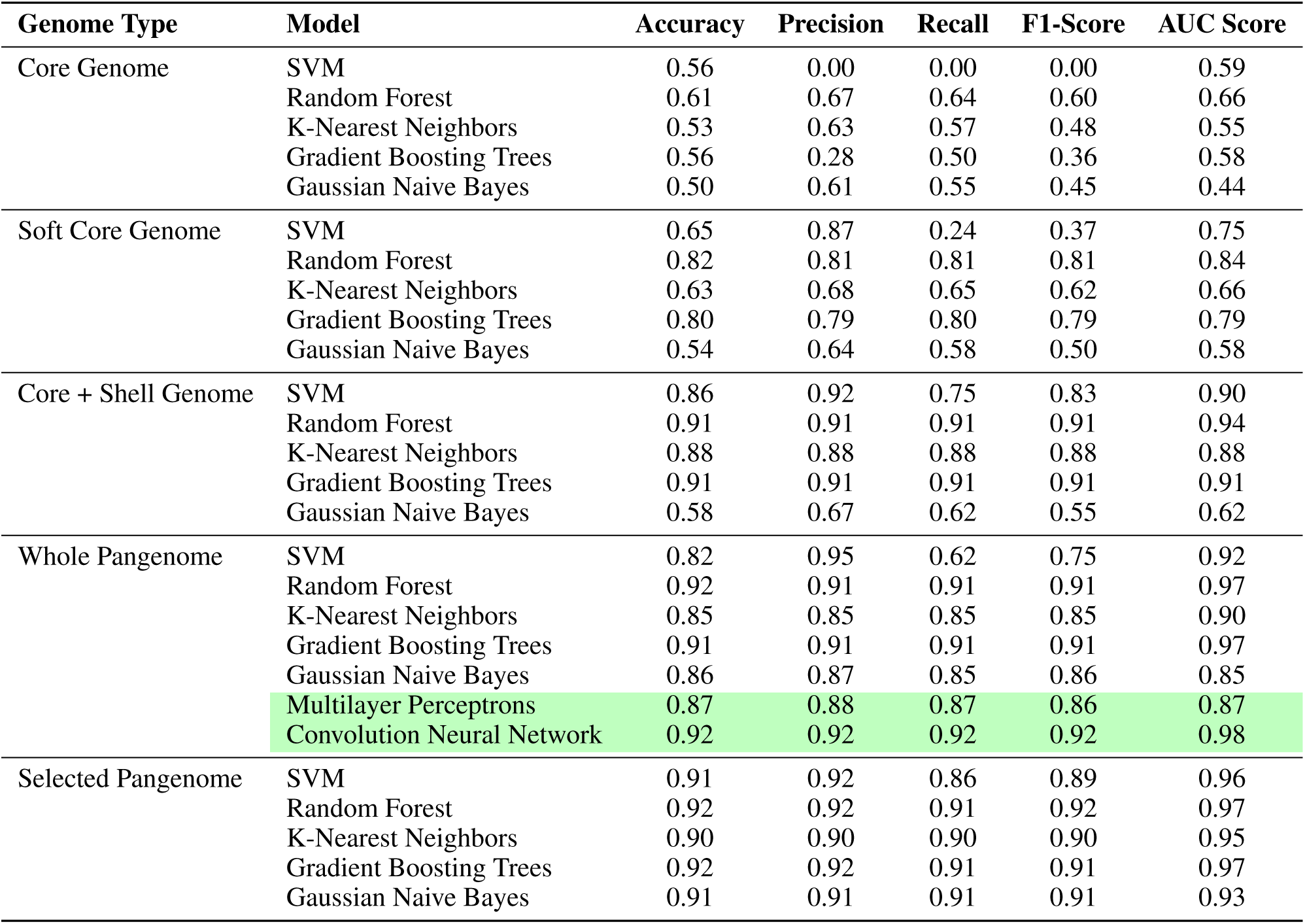
Supervised classification evaluation across different pangenome classes.

As shown in Fig 3, core genome provided limited useful signals, resulting in suboptimal performance for all models. Random Forest demonstrated the best performance among the classifiers, achieving an accuracy of 0.61 and an AUC score of 0.66. However, other models, such as SVM and K-Nearest Neighbors, struggled with low recall and F1-scores, reflecting their inability to determine pathogenic potential with this minimal feature set. The Gaussian Naive Bayes classifier, which assumes feature independence, had the lowest AUC score of 0.44, indicating that it struggled to capture the relationships within the core genome. These results highlight the limited discriminatory power of the core genome for classification tasks.

The addition of soft core genome expands the feature set by including genes shared among most, but not all, strains. This increase in genomic information led to substantial improvements in model performance. Random forest achieved the highest performance, with an accuracy of 0.82 and an AUC score of 0.84, showcasing its robustness to handle added soft core gene features. The SVM classifier also showed significant improvement, with an AUC score of 0.75, though its recall (0.24) and F1-score (0.37) remained relatively low, indicating its limitation to balance class predictions. Gradient Boosting Trees maintained stable performance with an AUC of 0.79, while Gaussian Naive Bayes still underperformed compared to other classifiers, with an AUC of 0.58. The results indicate that while the addition of s oft-core genomic features enhanced separability, models like SVM and Naive Bayes, still required additional optimization for effective classification.

The use of the core and shell genome provides a more comprehensive feature set by integrating accessory genes that are often associated with specific functions or adaptations. The core and shell combination showed a clear improvement in classification performance across all models. Random forest and Gradient Boosting Trees both achieved remarkable results, with accuracy and AUC scores reaching 0.91 and 0.94, respectively. SVM also showed strong performance, with an accuracy of 0.86 and an AUC of 0.90, demonstrating that it benefitted significantly from the additional features. K-Nearest Neighbors maintained high accuracy (0.88) but had slightly lower AUC scores compared to Random forest and Gradient Boosting Trees, suggesting a sensitivity to noise in the feature space. Gaussian Naive Bayes, while improved compared to the core genome, continued to lag behind other models, with an AUC of 0.62. These results underscore the utility of accessory genes in enhancing model performance and class separability.

The whole pangenome matrix incorporates the complete set of genes, including rare accessory genes, offering the most comprehensive genomic feature set. This led to the highest overall performance across all models. Random forest, Gradient Boosting Trees, and SVM each achieved exceptional accuracy (0.91–0.92) and AUC scores (0.92–0.97), demonstrating their ability to leverage the full range of genomic features for classification. K-Nearest Neighbors also performed well, with an accuracy of 0.85 and an AUC of 0.90, though slightly lower than the ensemble methods. Gaussian Naive Bayes improved significantly in this dataset, achieving an AUC of 0.85, indicating that it could better capture the relationships between features in this more extensive genomic context. These results highlight that the whole pangenome provides the richest feature set for classification, allowing models to achieve near-optimal performance.

### Feature selection was essential for ML model performance and training efficiency

Feature selection is crucial for enhancing both the performance and training efficiency of machine learning models, particularly for supervised classifiers (30). A prominent AUC score performance was illustrated in Fig 4. Across the progression from core genome to more feature-rich matrices such as soft core genome, core and shell genome, and whole pangenome, consistent improvements were observed across all models. Gaussian Naive Bayes, which initially struggled with limited features, showed significant boosts in accuracy and AUC scores as more genomic information was included, reaching its best performance with the selected pangenome (accuracy: 0.91, AUC:0.97) as shown in Fig4. Similarly, more complex models like Random Forest, SVM, and Gradient Boosting Trees exhibited substantial performance improvements as features increased. Random Forest consistently achieved strong results, with its AUC score improving from 0.66 on the core genome to 0.97 on the selected pangenome. SVM, which had low recall and F1-scores with limited features, leveraged the expanded feature sets to achieve a high AUC of 0.93 in the selected pangenome. Gradient Boosting Trees, known for its robustness to noise, also demonstrated near-optimal performance, with accuracy and AUC scores both reaching 0.92 and 0.97, respectively, on the selected pangenome.

**Figure 4:**
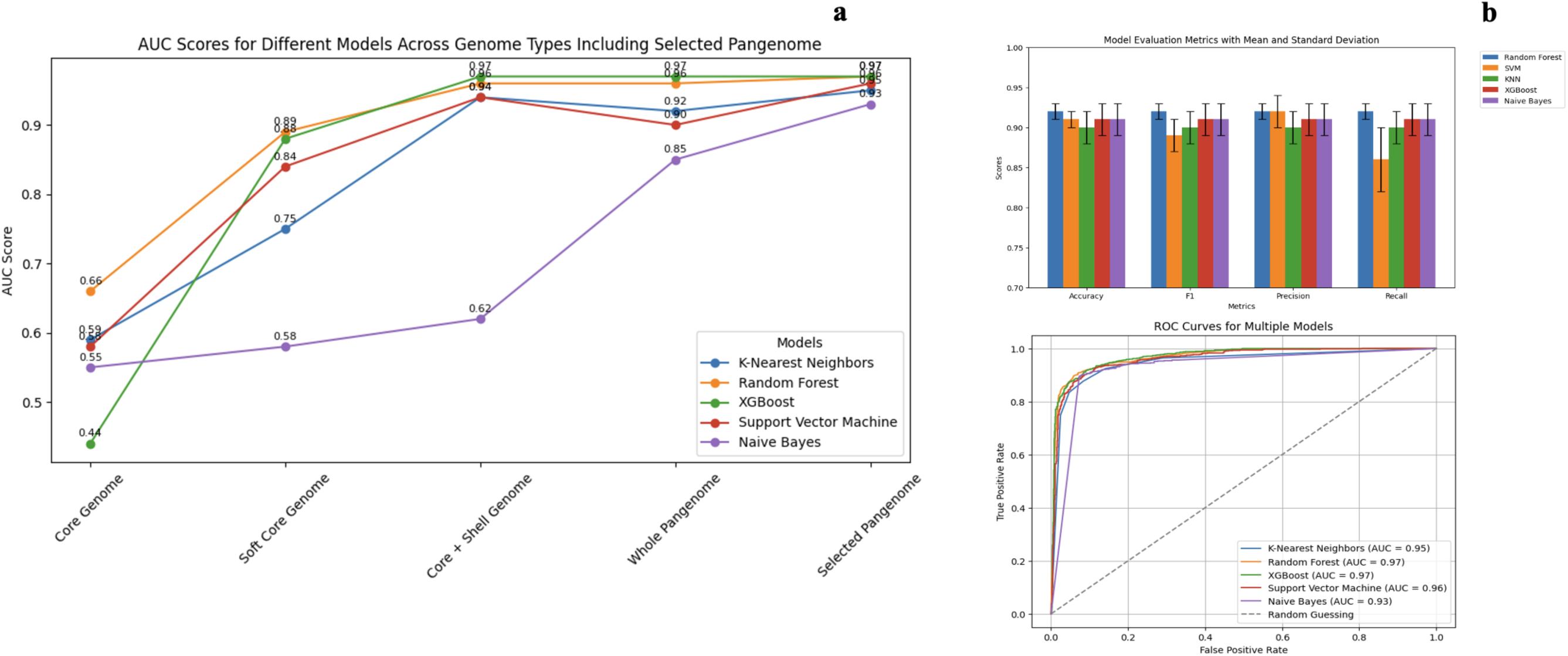
**a** AUC scores across different genome types for various models, including the selected pangenome features. The models compared are K-Nearest Neighbors, Random Forest, Gradient Boosting Trees, Support Vector Machine, and Naive Bayes, with AUC values annotated for each point. **b** Evaluation metrics (Accuracy, F1 Score, Precision, and Recall) with mean and standard deviation for all models across the selected genome features.

These results highlight that all models, regardless of complexity, benefit significantly from the inclusion of comprehensive and well-curated features. Feature selection not only boosts classification performance but also ensures efficient model training by focusing on the most relevant genomic information.

### Convolutional neural networks demonstrated high performance for classifying clinical and non-clinical strains using the pangenome matrix

In this study, the cloud genes within the pangenome play a significant role in pathogenic potential prediction proved by explainable models. Unlike core genes, which are conserved across strains, cloud genes are highly variable and are often associated with functions such as adaptation, pathogenicity, and resistance. These characteristics make it essential to analyze the entire pangenome, including the cloud genes, to fully capture the genetic variation that contributes to clinical relevance. Incorporating the entire pangenome ensures that potentially crucial information, particularly from cloud genes, is not overlooked during classification.

To evaluate supervised classification approaches, MLP was initially employed as they are commonly used for high-dimensional data. However, the results were suboptimal, as MLP struggled with the sparsity inherent in the pangenome matrix. This sparsity, characterized by the disproportionate distribution of non-zero entries across the matrix, diluted the contribution of cloud gene signals, resulting in limited performance. The inability of MLP to effectively isolate and amplify the localized signals from the cloud genes highlighted the need for a more sophisticated approach.

To address these challenges, convolutional neural networks (CNNs) were employed given their unique ability to extract spatial patterns and focus on localized regions of interest within high dimensional datasets. The CNN established in this study successfully conquered the matrix sparsity issue in cloud gene regions by concentrating the signal from specific regions through coevolutionary layers. This ability makes CNNs particularly suited for pangenome, where critical signals are scattered across the matrix in a sparse distribution. The decision to use CNNs for the high dimensional pangenome matrix was guided by the goal of fully utilizing all available genetic signals without relying on feature reduction techniques, which could risk losing important information.

As demonstrated in Table 3, CNNs outperformed other methods in classifying clinical and non-clinical strains, achieving an F1-Score and AUC score of 0.92 and 0.98, respectively. These results highlight the robustness of CNNs in handling complex, sparse datasets while preserving and amplifying meaningful signals. By enabling the comprehensive analysis of the pangenome, CNNs not only achieved superior classification performance but also underscored the importance of incorporating the entire genetic spectrum, including the highly variable cloud genes, in predictive models. This study illustrates the potential of CNNs as a powerful tool for genomic classification tasks, particularly when working with high-dimensional and sparse datasets like the pangenome matrix.

### Signals from non-core pangenome were key for pathogenic potential prediction as demonstrated by random forest feature importance and CNN grad-CAM analysis

To gain biological insights into the classification of clinical versus non-clinical strains, it is essential to employ interpretable models that can highlight the genomic features driving the predictions. Interpretability enables the identification of key genetic signals and provides a deeper understanding of the underlying biological mechanisms. To achieve this, two complementary approaches were utilized: Random Forest feature importance and Gradient-weighted Class Activation Mapping (Grad-CAM) for the convolutional neural network (CNN).

The analysis of selected gene features from the pangenome highlighted the critical role of non-core genes, particularly cloud genes, in differentiating clinical versus non-clinical strains. Figure 5a shows the feature importance rankings generated by the Random Forest model, where the top 1,980 gene features were identified based on their contribution to the classification efficiency. A significant proportion of these top-ranking features were classified as cloud genes, indicating their pivotal role in distinguishing strain types. This finding is supported by the distribution of pangenome levels in Figure 5b, where cloud genes constitute the majority (54.44%), followed by shell genes (43.74%) soft core genes (1.67%) and core genes (0.15%).

**Figure 5:**
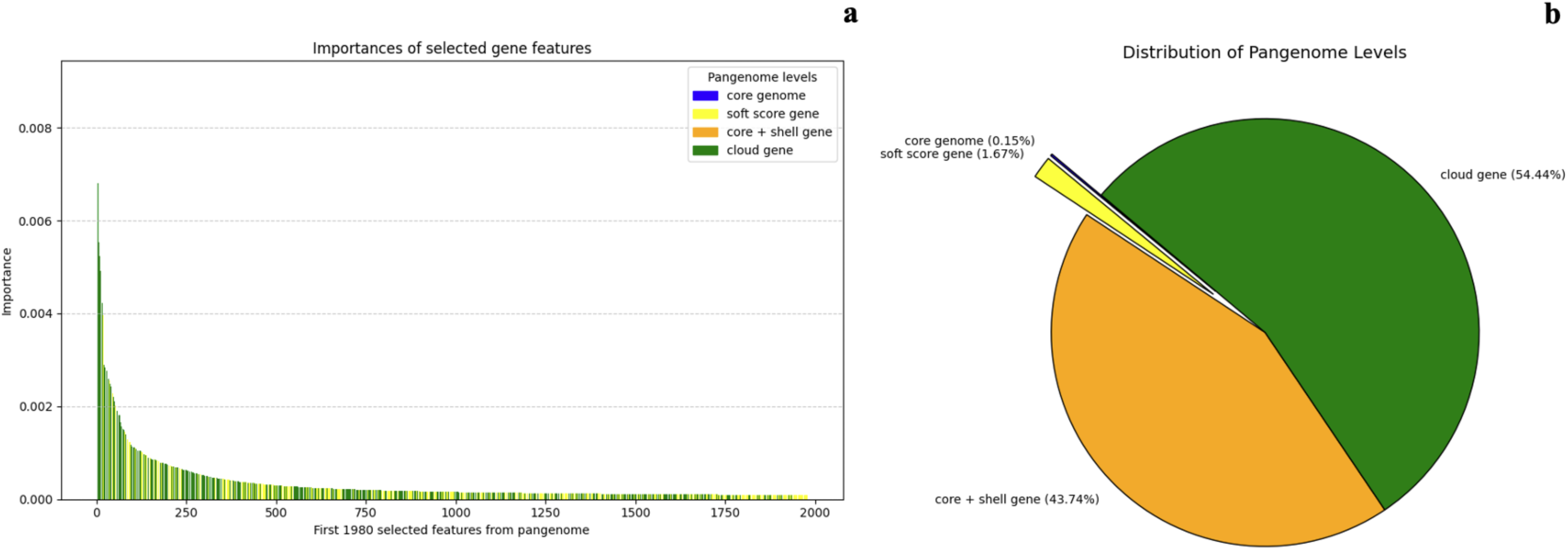
Overview of selected gene features from pangenome. (**a**) Importance of selected gene features: the importance weights of the top 1980 gene features as identified by a Random Forest model. (**b**) Distribution of pangenome levels:

These results underscore the limited contribution of core genes and emphasize the essential role of cloud genes in capturing strain-specific variations, particularly those associated with virulence and pathogenicity. The Random Forest model’s feature importance weights align closely with the genomic variability inherent in cloud genes, highlighting their significant impact on classification. Together, these findings demonstrate that the non-core pangenome, particularly cloud genes, is indispensable for effectively distinguishing clinical from non-clinical strains, supporting the need to consider the entire pangenome in classification models.

The Grad-CAM analysis of the CNN further corroborated the findings from the Random forest feature importance analysis, highlighting the critical role of non-core genes in classifying clinical versus non-clinical strains. As depicted in Figure 6, the heatmaps show the regions of the pangenome matrix where the model’s attention was most focused during classification. The areas of highest activation (indicated by warmer colors) align predominantly with the non-core pangenome regions, particularly the cloud genes.

**Figure 6:**
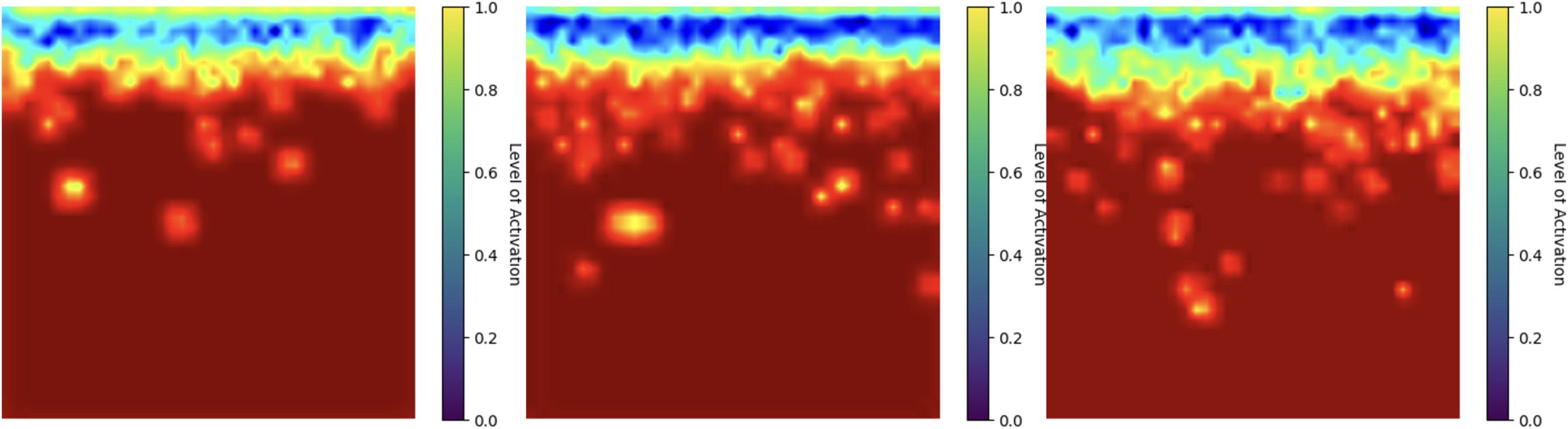
Three exemplary heatmaps depict the areas within the *Vp* whole-pangenome where model attention is focused, as determined by Gradient-weighted Class Activation Mapping (Grad-CAM) in CNN.

**Figure 7:**
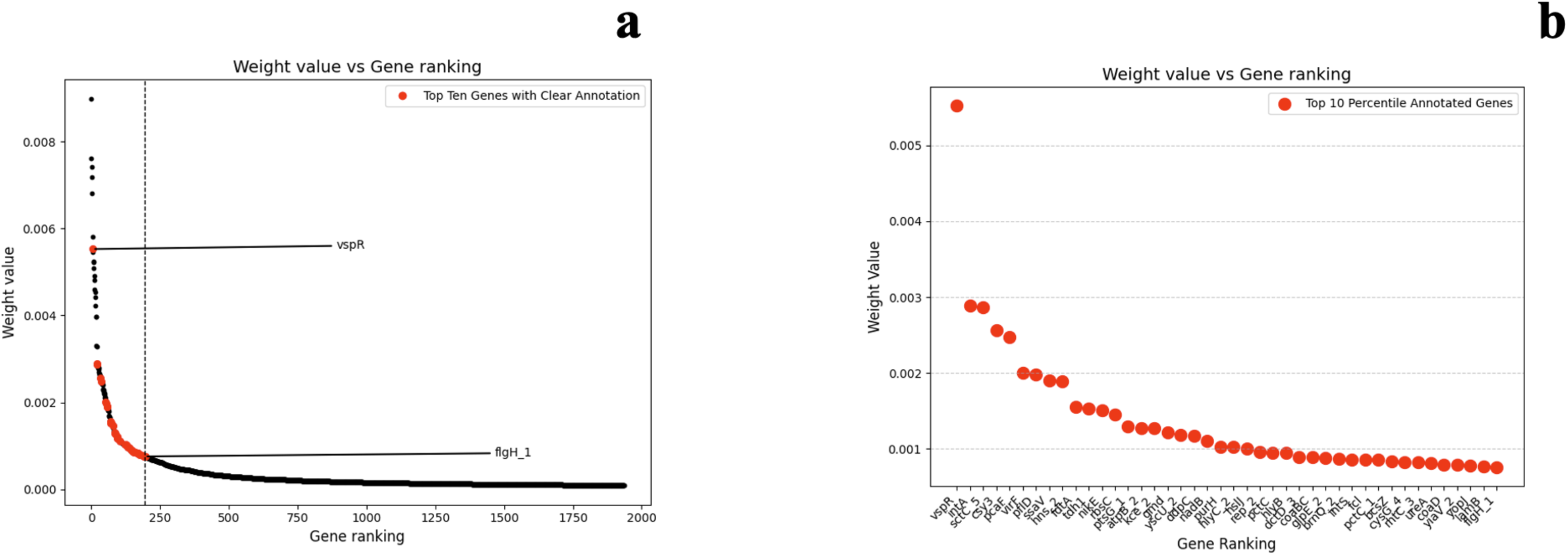
**(a)** Gene feature weight values assigned to genes by the Random Forest model across different rankings. The x-axis represents the gene ranking (ordered by weight value), and the y-axis shows the corresponding weight values. The red points highlight the genes in the top ten percentile with clear annotations from prokka. *vspR* and *flgH_1*, were annotated on the graph for reference the start and end annotated gene list respectively. **(b)** The zoomed-in view of the top 10 percentile of annotated genes from the ranking. The x-axis lists the gene names, and the y-axis displays their respective weight values.

This focus on non-core genomic features reinforced the importance of these highly variable genes in capturing strain-specific characteristics associated with pathogenic potential, which are often absent in the more conserved core genome. The alignment between the Grad-CAM results and the Random Forest feature importance weights emphasizes the consistent and indispensable contribution of non-core genes to the classification task, further validating their biological relevance and the necessity of considering the entire pangenome in predictive models.

## Discussion

Given its prevalence and negative impacts on both public health and the worldwide seafood industry, rapid and effective methods for characterizing *Vp* are imperatively needed to timely detect and identify *Vp* strains. By integrating pangenome-wise gene absence and presence patterns with advanced machine learning methods, this study addressed the imperative needs for rapid and accurate tools to differentiate clinical from non-clinical *Vp* strains. Throughout the evolutionary path of *Vp*, various genes associated with environmental adaptation have emerged, some of which might be associated with direct or indirect virulence-related properties (31). The latest comprehensive pangenomic analysis of *Vp* ST3 strains demonstrated how genetic divergence, selective pressures, and accessory gene acquisition during evolution facilitated its adaptation to the distinct local marine climate while also contributing to its virulence and ecological fitness (32). In another study, Jana, Keppel, Fridman, Bosis and Salomon (33) emphasized that acquired through horizontal gene transfer and localized within auxiliary modules or near DNA mobility elements, newly evolved virulence-associated genes in *Vp* pangenome are rare but essentially contributing its pathogenicity. To capture sparse but useful signals from the pangenomic matrix during the evolutionary trace of *Vp*, different layers of pangenome were developed and fit into learning models. Accordingly, results from this study demonstrated that incorporating more genes from the cloud gene component of the pangenome led to improved classification performance (Fig4), which underscored the significance of newly evolved genes within the *Vp* pangenome, particularly those localized in accessory genomic regions. Although unsupervised models were not able to predict pathogenic potential of *Vp* strains on high-dimensional pangenome matrix, supervised models demonstrated promising pathogenic potential prediction, especially for ensemble learning tree-based methods (AUC score 0.97 for Random Forest and Gradient Boosting Trees models). Slightly higher AUC score (0.98) was observed on CNN using full pangenome data input.

In the context of biological interpretability, Random Forest feature importance analysis provided deterministically ranked critical genes contributing the pathogenic potential of *Vp* whereas CNN Grad-CAM analyses highlighted the importance region of pangenome potentially associated with *Vp* virulence. Both approaches consistently identified non-core genes, particularly cloud genes, as the key drivers of classification. These genes are likely associated with strain-specific virulence and adaptability traits, making them prime targets for further functional and experimental validation. Given to the higher graduality of single gene level contribution to the virulence determination in Random forest compared with CNN (lost graduality due to gradient upscaling), the top 1936 genes with highest weight from Random forest was analyzed. Among these, *vspR*, a transcriptional regulator, emerged as a significant contributor. According to Davies, Bogard, Young and Mekalanos (34), VspR is a transcriptional inhibitor that represses the expression of the major pandemic island in *Vibrio cholerae* and the disruption of this gene recapitulate the intestinal colonization defect in TarB-mutant *V. cholerae*, which was supposed to fail in mouse intestine colonization. Meanwhile, in this study, *vspR* was ranked as the top one gene feature with the highest weight contributing the pathogencity determination of *Vp*. Consistent with substantial previous evidence, *tdh1* was found to play a significant role in determining the virulence of *Vp* further emphasizing the advantages of utilizing *tdh+* strains in research (12, 35–37). Additionally, several genes associated with the Type 3 Secretion System (T3SS) including sctC5 type III secretin and type III effector protein encoded gene yopJ were also highly ranked, suggesting the critical involvement of type III secretion system in the pathogenicity and host interaction mechanisms of *Vp*. Another top-ranked gene, *flgH1*, which encodes a structural component of the flagellar apparatus, underscored the importance of virulence factors associated with motility in colonization and persistence. These findings summarized in Table 4, highlight the pivotal role of specific genes in the pathogenicity of *Vp* and provide valuable insights for future research direction.

**Table 4:**
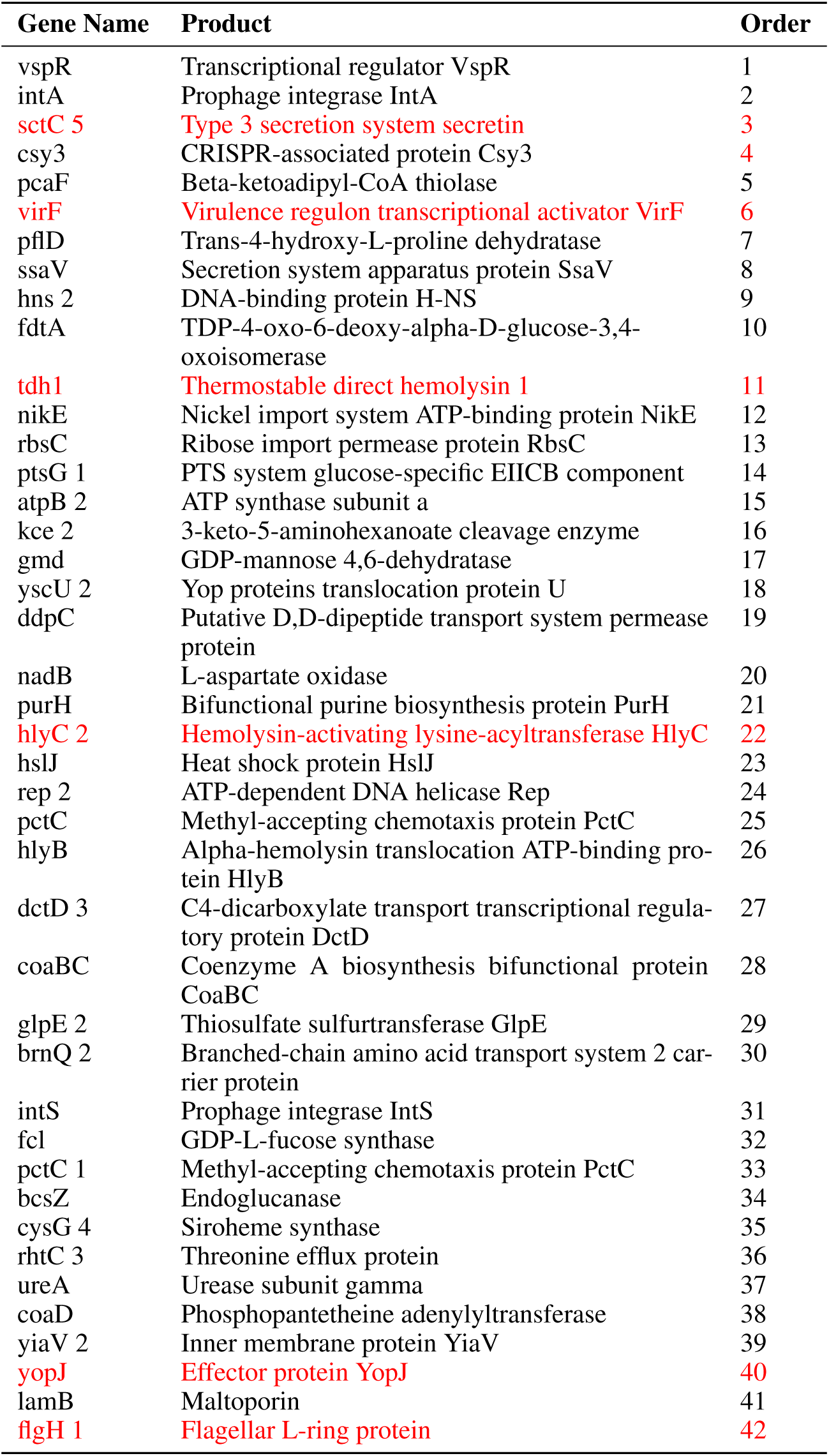
Ranked annotated genes contributing to *Vp* pathogenic potential prediction identified by Random Forest weight analysis (virulence-associated genes were highlighted in red).

It was noteworthy to point out that a significant proportion of virulence-associated genes in *Vp* remained functionally unknown (not being annotated by Prokka) and these genes usually do not serve as targets or usable references in conventional gene serotyping due to lack of sufficient information. Among the top 1936 genes contributing to its virulence, 1528 were categorized as functionally unknown (78.93%) (Fig7), emphasizing a large gap in gene annotation of their roles in pathogenicity. Notably, in the top 200 genes, 157 were also identified as functionally unknown (79.00%), further underscoring the need for focused research on these unexplored genes to uncover novel mechanisms driving virulence. These facts underscored the beauty of ML/DL models lies in their ability to leverage these functionally unknown genes to accurately predict the pathogenic potential of *Vp*, uncovering hidden patterns and relationships that traditional methods might overlook. The findings of this study provided critical insights into the application of machine learning approaches for the virulence characterization of *Vp* using whole-genome sequencing (WGS) data. By integrating pangenome-based features with advanced computational methods, this study addressed the pressing need for rapid and accurate tools to differentiate clinical from non-clinical *Vp* strains.

## Conclusion

By implementing ML and DL models based on pangenomic information of *Vp*, findings of this study have practical implications for public health and the seafood industry. The ability to rapidly and accurately predict the pathogenic potential of *Vp* strains can potentially inform risk assessments, outbreak investigations, and intervention strategies. Moreover, the integration of ML and DL models with WGS data aggregated to pangenomic data provides a scalable and reproducible framework for studying other foodborne pathogens such as *Listeria monocytogenes* and *Salmonella* spp (38, 39). This study underscores the transformative potential of ML and DL approaches in microbial genomics. By leveraging the full spectrum of the *Vp* pangenome, these methods enable accurate and interpretable predictions of pathogenicity, addressing a critical need in food safety research. The findings also highlight the importance of non-core genomic regions, particularly cloud genes, in capturing strain-specific traits. Together, these insights pave the way for the development of next-generation tools for pathogen characterization and control, contributing to improved public health outcomes and food industry practices.

## Data Availability

The data and code used for analysis are available in the GitHub repository at https://github.com/jlk666/VPML.

## Acknowledgment

We sincerely thank the Davis Data Lab at the University of California, Davis, for providing access to their NVIDIA A100 GPU, which significantly facilitated the training of our deep learning models.

